# A fluorescence correlation spectrometer for measurements in cuvettes

**DOI:** 10.1101/275230

**Authors:** B Sahoo, TB Sil, B Karmakar, K Garai

## Abstract

We have developed a fluorescence correlation spectroscopy (FCS) setup for performing single molecule measurements on samples inside regular cuvettes. We built this by using an Extra Long Working Distance (ELWD), 0.7 NA, air objective with working distance > 1.8 mm. We have achieved counts per molecule > 44 kHz, diffusion time < 64 μs for rhodamine B in aqueous buffer and a confocal volume < 2 fl. The cuvette-FCS can be used for measurements over a wide range of temperature that is beyond the range permitted in the microscope-based FCS. Finally, we demonstrate that cuvette-FCS can be coupled to automatic titrators to study urea dependent unfolding of proteins with unprecedented accuracy. The ease of use and compatibility with various accessories will enable applications of cuvette-FCS in the experiments that are regularly performed in fluorimeters but are generally avoided in microscope-based FCS.

---

Fluorescence correlation spectroscopy (FCS) is a powerful single molecule technique with wide spread applications in biophysics and biology. The high spatio-temporal resolution of FCS allows measurements of fast processes in dilute solutions. Some of the common applications of FCS are measurements of molecular size, chemical kinetics, conformational dynamics of biomolecules, protein-ligand interactions and protein aggregation in vitro (1-4). Due to its single molecule sensitivity, FCS techniques have been developed extensively over the past two decades for measurements in live cells and tissue samples. For example, several flavors of imaging FCS techniques have been developed for measurements of membrane dynamics, cellular and nuclear localization, and transport of biomolecules in live cells (5-8). Furthermore, development of 2-photon FCS has extended applications of FCS in tissues and other thick samples (9).

Conventionally, FCS setups are attached to confocal microscopes for the high spatial resolution and the signal to noise (S/N) achieved in confocal microscopy. Consequently, applications of FCS have several limitations in many of the biophysical experiments. For example, folding-unfolding of proteins using chemical or thermal denaturation are rarely studied using FCS (10-12). Kinetic experiments which require stirring of the samples can’t be performed using microscope based FCS setups. Furthermore, experiments that require non-aqueous or corrosive solvents are seldom performed by using FCS (13). However, the above mentioned experiments are performed regularly in most biophysics and biochemistry laboratories using fluorimeters. Therefore, an FCS setup capable of performing measurements inside a cuvette with high S/N can enable applications of FCS in the above-mentioned biophysical experiments conveniently.

A major obstacle for performing FCS in a cuvette is that the thickness of the optical windows of commercially available cuvettes are about 1.25 mm. Conventional FCS setups employ high numerical aperature (NA ≥ 1.2) objective lenses, which are carefully corrected for both spherical and chromatic aberrations. These objectives have working distances less than 0.31 mm, and hence can’t be used for measurements inside cuvettes. While high NA objectives are required for high S/N, FCS measurements have been demonstrated using optics with much lower NA. For example, Garai et al performed FCS measurements using a single mode optical fiber (NA = 0.13) for detection of protein aggregates in samples placed remotely (14). Furthermore, Banachowicz et al. have shown that FCS measurements can be performed using objectives with NA equal to 0.4 (15).

Here we report building a highly senstitive FCS setup capable of performing measurements inside regular cuvettes. This setup is built using an extra long working distance (ELWD) objective can yield molecular brightness > 44 kHz and a diffusion time (τ_D_) < 64 μs for Rhodamine B in phosphate buffered saline (PBS) at pH 7.4. The confocal volume obtained in this setup is < 2 fl.

Figure 1 shows the schematic of the cuvette-FCS setup. This setup is similar to the conventional FCS with two major differences. First, the objective is mounted horizontally. Such geometry is required for FCS measurements inside a cuvette. Second, we have used an ELWD plan achromat objective with NA equal to 0.7. These objectives are corrected for both spherical and chromatic aberrations but the corrections are less extensive than the conventionally used plan apochromat objectives. However, the advantage of using the ELWD objective is that it’s working distance is > 1.8 mm and the correction collar can be adjusted for cover slips of thickness up to 1.3 mm. Therefore, these objectives may be suitable for use with common cuvettes.

**Figure 1:**
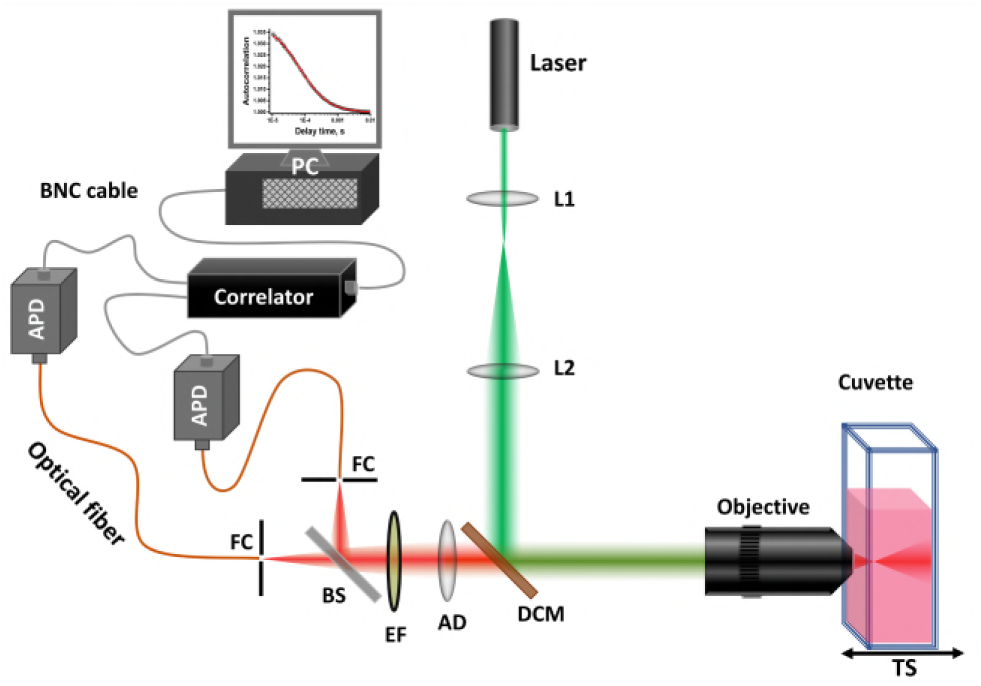
Schematic of the cuvette-FCS setup. Laser (543 nm, CW), L1 and L2: lenses, DCM: Dichroic mirror, AD: Achromatic doublet lens, EF: emission filter, BS: 50/50 beam splitter, FC: fiber coupler, TS: micrometer translation stage, objective: ELWD, air objective, 0.7 NA, WD = 2.6-1.8 mm.

We have used a solution of rhodamine B in PBS to characterize the sensitivity and the spatial resolution of the cuvette-FCS setup. Supplementary Eq. S1 shows that FCS autocorrelation (G(τ)) data can be analyzed to determine the average number of the molecules (<N>) and the diffusion time (τ_D_) of the fluorophore in the FCS observation volume. The confocal volume (Vconfocal) and the axial resolution (σ_xy_) can be determined from <N> and τ_D_ respectively (Supplementary Eq. S2 and S3). Furthermore, S/N of the G(τ) is dependent on the number of photons collected per molecule per unit time, viz, the counts per molecule (CPM) (16). CPM is calculated from the ratio of total photon count rate and <N>.

First, we investigate the factors that affect the performance of the cuvette-FCS most ctirically. We find that τ_D_, <N> and CPM obtained are critically dependent on two factors, viz, quality of the cuvettes, and alignment of the cuvette with respect to the objective. Figure 2A compares the autocorrelation data obtained from an aqueous solution of rhodamine B placed in quartz cuvettes procured from three different manufacturers referred here as Hellma, MF1 and MF2. Dimensions of all the three cuvettes are the same, e.g., thickness of the optical window = 1.25 mm and the optical pathlength = 10 mm. Amplitudes of the autocorrelation traces (i.e., G(0)) obtained from the different cuvettes differ by about 1.4-fold. Furthermore, supplementary Figure S2A and S2B show that τ_D_ and CPM differ by 1.2 and 2.0-fold respectively. Therefore quality of the optical windows of the cuvettes is extremely important for the S/N of the FCS data. It is clear that the Hellma cuvette is the most suitable for cuvette-FCS. Hence, all the FCS measurements reported in the rest of this article are performed using a Hellma cuvette.

**Figure 2:**
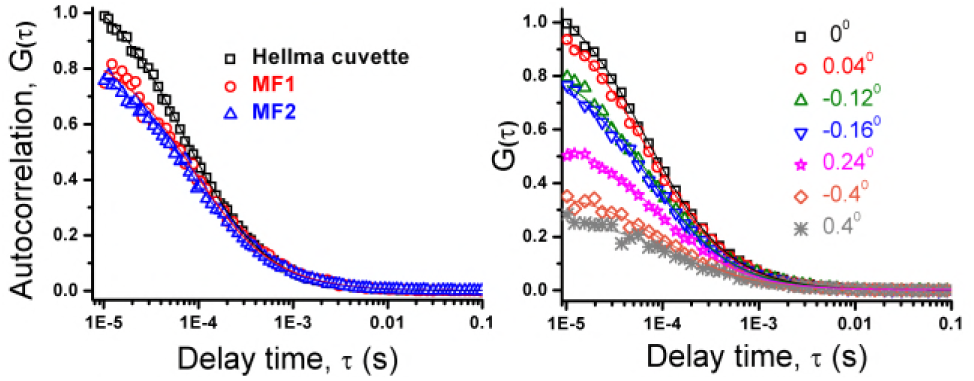
Effects of quality and angular misalignment (θ) of the cuvette on the autocorrelation, G(τ) data. A) G(τ) obtained from aqueous solution of rhodamine B using cuvettes from three different manufacturers. G(τ) using Hellma cuvette is the best. B) G(τ) at different θ. G(τ) worsens drastically with even a small (by ±0.40^0^) increase of θ. Hence, the cuvette must be placed parallel to the objective. The symbols represent data and the solid lines are fits using single diffusing component (suppl. Eq. S1).

We then examined the effects of the relative angle between the face of the objective and the optical window of the cuvette. Figure 2B shows that G(τ) changes drastically with the change of the angle. It may be seen that G(0) decreases by 4 fold for an angular misalignment of 0.40 degree. Supplementary Figure S3A and B show that corresponding increase in τ_D_ and decrease in CPM are about 2 and 4-folds respectively. Hence, the best sensitivity in the cuvette-FCS can be achieved when the cuvette surface is parallel to the face of the objective.

We then calibrate the cuvette-FCS setup by measuring the CPM, <N> and τ_D_ using solutions of rhodamine B in PBS buffer at pH 7.4. Figure 3A shows that CPM of rhodamine B increases monotonically with the incident laser power. The increase of CPM deviates from linearity above 50 µW of incident power but it doesn’t reach complete saturatation even at the highest intensity (114 μW) used here. It may be seen that the maximum CPM obtained here from rhodamine B is 44 kHz. Supplementary Figure S4B shows that average τ_D_ obtained from fitting all the G(τ) data is 64±2 μs.

**Figure 3:**
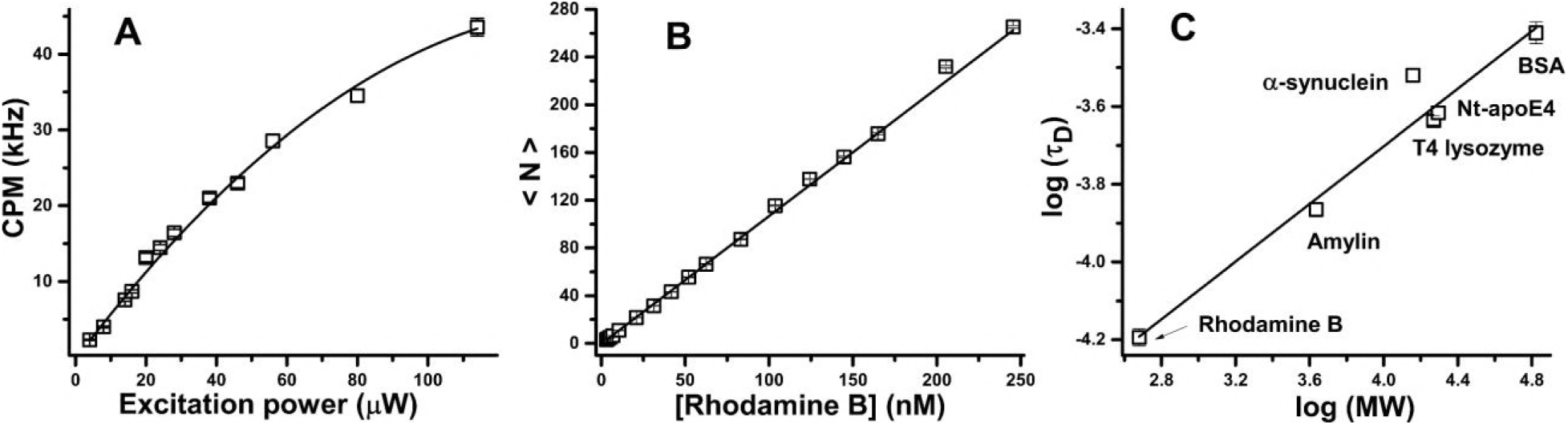
Characterization of sensitivity and resolution of the cuvette-FCS Setup. A) CPM as a function of incident laser power obtained using aqueous solution of rhodamine B. Symbols represent data and the solid line is fit using a parabola. CPM increases linearly till ≈ 50 µW of incident power. B) Average number of molecules (<N>) in the confocal observation volume (*V*_*confocal*_) as a function of concentration of rhodamine B. Symbols represent data. Slope of the fitted line is 1.1 molecules/nM indicating *V*_*confocal*_ = 1.8 fl. C) log-log plot of τ_D_ versus molecular weight (MW) of free dye and several TMR labeled-proteins. Symbols represent data and the solid line is linear fit. The slope obtained is 0.36±0.01. Expected slope for globular proteins is 0.33 (17).

Figure 3B shows the plot of *<N>* as a function of concentration (*C*) of rhodamine B. As expected, *<N>* varies linearly with *C* over the entire range of concentration of 3-265 nM. Linear fit of the data yields a slope = 1.1 molecule/nM. Hence, the estimated value of *V_confocal_* in cuvette-FCS is 1.8 fl (see Supplementary Eq. S2). Supplementary Figure S5B shows that mean τ_D_ of rhodamine B obtained from fitting of all the G(τ) data is 63±2 μs. Using the known diffusion coefficient (D = 4.5×10-6 cm^2^/s) of rhodamine B in water, the axial resolution in cuvette-FCS may be estimated as 340 nm (see supplementary Eq. S3). The axial resolution expected for a diffraction limited confocal setup is between 285 to 422 nm for *λ*_ex_ = 543 nm and NA = 0.7 (see supplementary Eq. S4). Hence, axial resolution of the cuvette-FCS setup is within the expected range.

We then examine if cuvette-FCS is suitable for measurements using proteins. The proteins used here are amylin, α-Synuclein, T4-Lysozyme, an N-terminal fragment of apolipoprotein E4 (Nt-ApoE4) and bovine serum albumin (BSA). All the proteins are fluorescently labeled with tetramethylrhodamine (TMR). Figure 3C shows that log(τ_D_) varies linearly with log(MW). The slope of the line is equal to 0.36 ± 0.01. The expected scaling exponent is 0.33 for globular proteins (17). Hence our data are consistent with the scaling law for globular proteins. The τ_D_ of α-synuclein is found to be somewhat higher, which is consistent with the extended conformation of α-synuclein in native conditions (18).

We then examine suitability of the cuvette-FCS setup for measurements in conditions that are generally avoided in microscope based FCS. First, we perform FCS measurements using rhodamine B in a range of temperature starting from 15° to 60°C. In confocal microscopes temperature range is generally restricted to 25°-37°C due to incompatibility of the objectives at the higher or the lower temperatures. Since we are using an air objective the temperature of the sample doesn’t affect the resolution. As viscosity of water changes with temperature we have plotted τ_D_ of rhodamine B as a function of viscosity of the solution in Figure 4A. It may be seen that τ_D_ varies linearly with the viscosity of water as may be expected from Stokes-Einstein (SE) relationship (19). Hence, Cuvette-FCS is suitable for measurements over a wide range of temperature. While here we have used a limited range of temperature, with appropriately designed cell holder even larger range of temperature can be used for measurements using cuvette-FCS.

**Figure 4:**
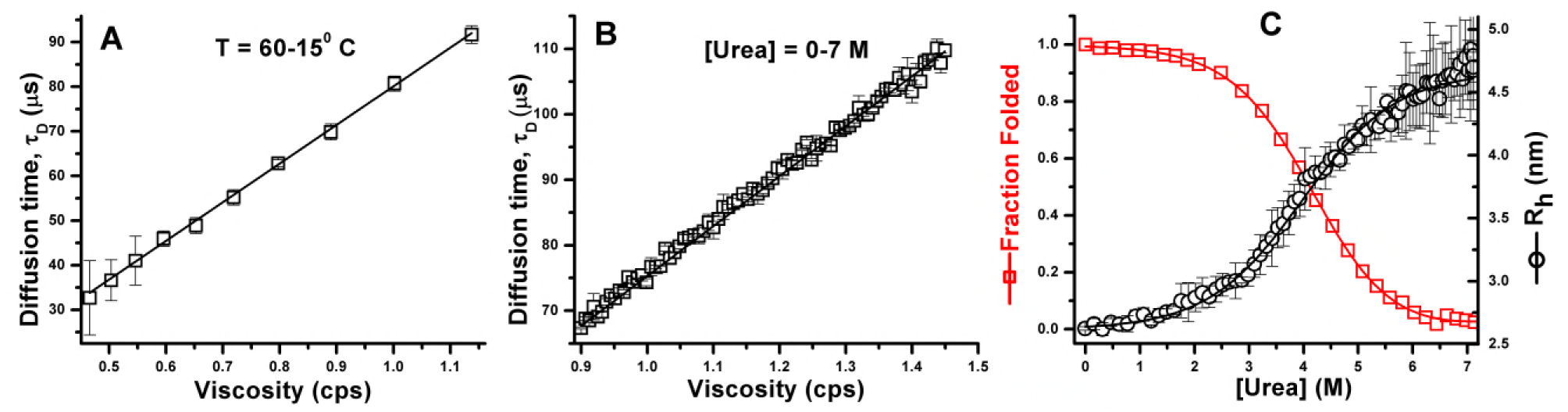
Applications of cuvette-FCS in a range of temperature (A) and in urea (B). A) τ_D_ of rhodamine B versus viscosity. Here visocity changes due to change of temperature of the solution. (B) τ_D_ versus viscosity of urea. Symbols represent data and the solid line is linear fit of the data. Both A and B show that τ_D_ increases linearly with viscosity, consistent with the Stokes-Einstein relationship (11). (C) Urea dependent unfolding of TMR-Nt-ApoE4. The circles represent hydrodynamic radius (R_h_) measured by FCS and the squares represent fraction folded measured by CD at 222 nm. The solid lines are fits using a two-state model (21). Clearly, increase of R_h_ is consistent with loss of secondary structure due to unfolding of the protein

We then examine if the cuvette-FCS can be used for measurements in urea. Urea is widely used as a chemical denaturant in experiments involving folding-unfolding of proteins. While FCS can yield single molecule level information, there are major difficulties in measurements of molecular size in urea in conventional FCS setups. The difficulties arise due to mismatch of refractive index of the solution, requiring adjustment of the distance between the objective and the glass coverslip for samples containing urea. These complications have been discussed in detail elsewhere (10). Cuvette-FCS has certain advantages in these experiments. For example, the distance between the objective and the cuvette is fixed during the entire duration of the experiment. Concentration of urea can be changed by manual pipetting or by using autotitrators as has been done here. Furthemore, mixing of the samples can be performed by stirring using magnetic stirrers. Therefore, titration experiments in the cuvette-FCS setup can be automated fully. Figure 4B shows the effect of concentration of urea on the τ_D_ of rhodamine B. These experiments are performed using an automatic titrator coupled to the cuvette, which is placed in a temperature controlled cell holder. The entire experiment is performed in an automatic manner. It may be seen that the τ_D_ of rhodamine B increases linearly with increase of viscosity of the urea solution as expected from SE relationship shown earlier by others (11). We note here that in this experiment FCS measurements have been performed on a total of 77 intermediate concentrations of urea between 0-7.1 M. Measurements using such large number of concentrations of urea are possible in this setup due to compatibility of cuvette-FCS with automatic titrators.

Finally, we examine if cuvette-FCS can be successfully employed to study unfolding of proteins induced by urea. Here we have used a folded protein, viz, TMR-labeled-Nt-apoE4). The Nt-apoE4 is a four helix bundle (PDB ID, 1gs9). The hydrodynamic radius (Rh) of Nt-apoE4 calculated using Hydropro10 is 2.67 nm (20). Figure 4C shows that R_h_ of Nt-apoE4 increases monotonically from 2.6 nm to 4.8 nm with increasing concentrations of urea from 0 to 7.1 M. The R_h_ of Nt-apoE4 shows a sigmoidal transition consistent with cooperative unfolding of the protein in urea. Furthermore, urea dependent changes of R_h_ agree well with the changes in secondary structure content measured by Circular Dichroism (CD). The midpoint of transition measured by FCS and CD respectively are 4.0 and 3.9 M of urea when fitted with a two-state model (21). However, FCS data show that R_h_ continues to increase even after complete unfolding of the protein, indicating expansion of the unfolded chain at higher concentrations of the denaturant (11, 12, 22).

Here we demonstrate that FCS measurements can be performed inside regular cuvettes with high sensitivity. In our current setup we have obtained CPM greater than 44 kHz from aqueous solution of rhodamine B and a confocal volume less than 2 fl. Expectedly, the sensitivity and the spatial resolution are lower than that can be achieved in our homebuilt microscope-based FCS setup (data not shown). However, the maximum CPM obtained using commcercial microscope based FCS are reported to be about 30 – 50 kHz (from alexa488 in aqueous buffer) and the confocal volume obtained is about 1.0 fl (11, 23). Therefore, the sensitivity and the spatial resolution of cuvette-FCS is comparable to some of the commercially available microscope-based FCS.

Cuvette-FCS offers several advantages over microscope based FCS instruments. Two major advantages have been demonstrated here. For example cuvette FCS allows measurements over a large range of temperature. Furthermore, cuvette-FCS can be integrated with automatic titrators for fully automated titration experiments. Use of automatic titrators and temperature controlled cell holders are particularly advantageous in acquiring large number of highly accurate data points as may be seen in Figure 4B and C. Sherman and Haran have used single molecule FRET and FCS to measure GdnCl induced expansion of protein L to determine the coil-globule (CG) transition point and calculated the per residue average solvation energy (19). Furhermore, Schuler and coworkers have demonstrated that single molecule studies of protein unfolding can be used to estimate net intrachain interaction energy of the unfolded form of a protein in a particular solvent (12). However, there are only a few studies of protein unfolding using FCS reported in the literature (10, 11, 19). We have demonstrated that Cuvette-FCS can be used to study chain collapse or expansion respectively due to folding or unfolding of proteins with high sensitivity and accuracy (see figure 4C). Protein unfolding data reported here show that changes in secondary structure and the hydrodynamic size of the Nt-apoE4 are almost concomitant. These results are consistent with those reported earlier for denaturation of IFABP (10). However, Sherman et al have found that the midpoints of transition measured by FCS and by CD are somewhat different in case of adenylate kinase (AK) protein (11). We speculate here that cuvette-FCS can be useful in addressing several other problems. For example, FCS has been used to monitor formation of oligomers and protofibrils in the early stages of protein aggregation (24). However, nucleation dependent aggregation kinetics are generally slow, occuring over several hours, days or weeks. Therefore, in vitro these processes are accelarated by agitations such as stirring and/or by raising the temperature of incubation (25). Cuvette-FCS would be suitable to monitor such processes. Cuvette-FCS can also be used for measurements in non-aqueous solvents including corrosive solvents such as those experiments involving solvent dependent dynamics of colloids, polymers or biopolymers such as intrinsically disordered proteins. Furthermore, cuvette-FCS can offer significant advantage in measurements at very low concentrations of biomolecules due to the low surface to volume ratio of cuevttes and due to compatibility of its surfaces to passivation by covalent linkage of PEG to minimize adsorption (26).

Finally, we propose that our setup can be integrated with fluorimeters. Fluorimeters are highly popular for applications in wide range of experiments in biochemistry and biophysics laboratories. However, fluorimeter measurements provide only ensemble level information. Therefore, cuvette-FCS can be extremely useful in performing single molecule measurements in most of the experiments that are performed regularly in fluorimeters but not performed in microscope-based FCS instruments.

**Author Contributions**

KG conceptualized the experiments. BS, TBS, BK and KG designed and performed research. BS and KG analyzed data and wrote the paper.

## Acknowledgements

We acknowledge funding from Tata Insititute of Fundamental Research.

